# Ultra-fast micro-CT of an unrestrained live insect

**DOI:** 10.1101/2023.03.03.531017

**Authors:** Leonidas-Romanos Davranoglou, Beth Mortimer, Christian Matthias Schlepütz, Graham K. Taylor

## Abstract

Micro-CT has revolutionized functional morphology by enabling volumetric reconstruction of biological specimens at micrometre scales, but its accuracy is compromised by fixation artefacts. State-of-the-art *in vivo* imaging avoids this, but still requires subjects to be tethered, anaesthetized, or stained. Here we use ultra-fast synchrotron-based micro-CT to produce the first 3D scan of an unrestrained living organism at micrometre resolution, demonstrating the potential of this method in physiology, behaviour, and biomechanics.

---

X-ray micro-computed tomography (micro-CT) is a key method in functional morphology, enabling the non-destructive 3D imaging of microscopic internal and external structures that would otherwise be impossible to study^1–3^. Conventional micro-CT requires specimens to be fixed to avoid movement artefacts, but the fixation artefacts this introduces (e.g., desiccation, or shrinkage of specimens in ethanol) impedes the accuracy of quantitative anatomical measurements^1,4,5^. Fixation artefacts can be eliminated completely by using fast *in vivo* imaging methods, which to date has enabled cineradiographic imaging^5,10^ of live specimens and even tomographic reconstruction^3,6–9^ of periodic 3D motions. Nevertheless, subjects have had to be tethered^5,6,10^, sedated^9^, or stained^9^ before scanning, which introduces other behavioural and anatomical artefacts. Here we use ultrafast, synchrotron-based micro-CT to generate what is, to our knowledge, the first volumetric image of an unrestrained living organism at micrometre resolution.

We allowed a small (∼4.5 mm body length) planthopper insect (*Phantia subquadrata*, Hemiptera) to walk freely on a plant stem attached to a rotation stage within the field-of-view of the TOMCAT beamline at the Swiss Light Source, Paul Scherrer Institut, Switzerland (Figure 1A). Tomographic reconstruction requires the collection of a consistent set of radiographs as the specimen is rotated through at least one 180° half-scan, but the quality of the reconstruction can be improved by collecting one or more 360° scans. We set the stage rotating at an angular speed 720° s^−1^ and allowed the insect to acclimatize with the beam shutter closed. Once the insect had settled, we initiated high-speed imaging, collecting a sequence of 1598 radiographs at a pixel size of 4.9 × 4.9 μm and a sampling rate of 2 kHz. We used a detection system comprising a 100 μm thick LuAG:Ce scintillator, an optical microscope operating at 2.24 times magnification, and a GigaFRoST camera^11^ recording images at a 0.5 ms exposure time, with a beam energy of 16 keV.

**Figure 1.**
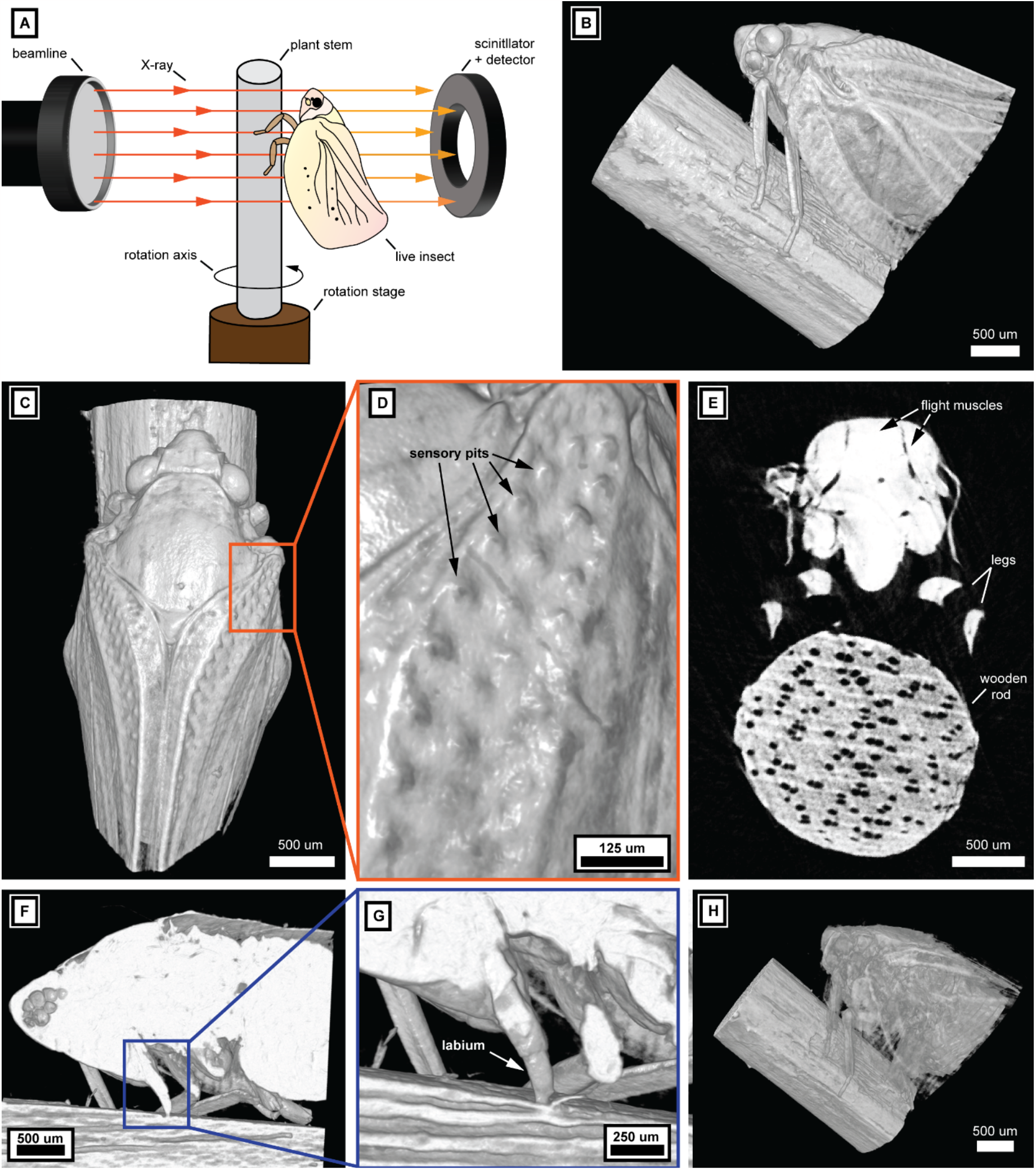
High-speed synchrotron-based X-ray microtomography of an unrestrained live insect. **(A)** Schematic of experimental setup. **(B-D)** 3D volumetric reconstruction of living planthopper *Ph. subquadrata* during the first half-scan, viewed from the side (B) and dorsally (C), with magnified inset (D) showing the sensory pits on the right forewing. **(E)** Cross-section of tomogram at mid-mesothorax, showing a portion of the first set of wing muscles. **(F)** Lateral-longitudinal section of tomogram, with blue inset focusing on the labium. **(G)** Closeup of labium probing the surface of the stem, interpreted as an attempt at feeding. **(H)** 3D volumetric reconstruction of the second half-scan, during which the insect moved, resulting in a recognizable but motion-blurred image.

The insect remained stationary throughout the entire 250 ms duration of the first half-scan. This enabled a high-resolution 3D volumetric reconstruction of its morphology using phase retrieval, at an isotropic voxel spacing of 4.9 μm (Figure 1B-D, F, G). The insect began walking up the stem during the second half-scan, as can be seen in the cineradiographic projections. This resulted in a motion-blurred 3D reconstruction, from which only limited information could be recovered (Figure 1H). The 3D volumetric reconstruction acquired during the first half-scan captures the external morphology of the insect in exquisite detail (Figure 1B-D). Microscopic sense organs called sensory pits^12^, are clearly visible on the wings, measuring only tens of micrometres across (Figure 1D). The insect also appears to have been attempting to feed at the start of the scan, because its syringe-like mouthparts (labium) can be seen probing the surface of the plant stem (Figure 1F,G). Such behavior would be challenging or impossible to capture with the slow scan times available at most current micro-CT facilities. The internal structure of the insect was poorly resolved (Figure 1E), because only one half-scan could be completed before the insect began moving. Only the principal flight muscles are visible (Figure 1H), distinguished by virtue of the air-filled cavities between them. In contrast, the internal microstructure of the plant stem was adequately reconstructed (Figure 1E), probably owing to its lower material density.

Overall, our study demonstrates that ultra-fast synchrotron X-ray microtomography can be used to create 3D reconstructions of unrestrained, living arthropods, greatly extending the potential of this technique in animal physiology, behaviour, and biomechanics. Possible applications of this methodology include recording the 3D pose of living organisms at static equilibrium (e.g. during stationary behaviors including feeding and ovipositing on plants or inside other organisms in the case of parasitoids), the measurement of growth rates of different life stages that cannot be observed externally (e.g. larvae of wood-boring insects, developing embryos in medical pests such as tsetse flies), and visualization of internal and external structures without the risk of preservation artefacts.

## Acknowledgments

L-R.D. is a Leverhulme Trust Early Career Research Fellow and John Fell OUP Fellow, and was supported by an Oxford-NaturalMotion scholarship and the Alexander S. Onassis Public Benefit Foundation Scholarships for Hellenes during this work. B.M. is a Royal Society University Research Fellow, and was supported by a Royal Commission for the Exhibition of 1851 Research Fellowship during this work. G.T. was supported in this work by a grant from Jesus College, Oxford. We acknowledge the Paul Scherrer Institut, Villigen, Switzerland for provision of synchrotron radiation beamtime at the TOMCAT beamline X02DA of the SLS.

## Author contributions

Conceptualization: L-R.D, C.M.S., G.T., B.M. Data curation: L-R.D., C. M.S. Formal Analysis: L-R.D, C. M.S.,

G.T., B.M. Funding acquisition: L-R.D,, G.T., B.M. Investigation: L-R.D, C. M.S., G.T., B.M. Methodology: L-R.D, C. M.S., G.T., B.M. Supervision: G.T., B.M. Visualization: L-R.D. Writing – original draft: L-R.D. Writing – review and editing. G.T.

## Notes

### Competing Interest Statement

The authors have declared no competing interest.

